# FAVOR: Functional Annotation of Variants Online Resource and Annotator for Variation across the Human Genome

**DOI:** 10.1101/2022.08.28.505582

**Authors:** Hufeng Zhou, Theodore Arapoglou, Xihao Li, Zilin Li, Xiuwen Zheng, Jill Moore, Abhijith Asok, Sushant Kumar, Elizabeth E. Blue, Steven Buyske, Nancy Cox, Adam Felsenfeld, Mark Gerstein, Eimear Kenny, Bingshan Li, Tara Matise, Anthony Philippakis, Heidi Rehm, Heidi J. Sofia, Grace Snyder, NHGRI Genome Sequencing Program Variant Functional Annotation Working Group, Zhiping Weng, Benjamin Neale, Shamil R. Sunyaev, Xihong Lin

## Abstract

Large-scale whole genome sequencing (WGS) studies and biobanks are rapidly generating a multitude of coding and non-coding variants. They provide an unprecedented resource for illuminating the genetic basis of human diseases. Variant functional annotations play a critical role in WGS analysis, result interpretation, and prioritization of disease- or trait-associated causal variants. Existing functional annotation databases have limited scope to perform online queries or are unable to functionally annotate the genotype data of large WGS studies and biobanks for downstream analysis. We develop the Functional Annotation of Variants Online Resources (FAVOR) to meet these pressing needs. FAVOR provides a comprehensive online multi-faceted portal with summarization and visualization of all possible 9 billion single nucleotide variants (SNVs) across the genome, and allows for rapid variant-, gene-, and region-level online queries. It integrates variant functional information from multiple sources to describe the functional characteristics of variants and facilitates prioritizing plausible causal variants influencing human phenotypes. Furthermore, a scalable annotation tool, FAVORannotator, is provided for functionally annotating and efficiently storing the genotype and variant functional annotation data of a large-scale sequencing study in an annotated GDS file format to facilitate downstream analysis. FAVOR and FAVORannotator are available at *https://favor.genohub.org*.

## INTRODUCTION

A rapidly increasing number of large-scale Whole Genome/Exome Sequencing (WGS/WES) studies and biobanks are being conducted. They provide rich opportunities for understanding the genetic bases of complex human diseases and traits. Examples of large WGS/WES studies and biobanks include the Trans-Omics Precision Medicine Program (TOPMed) of the National Heart, Lung and Blood Institute (NHLBI) (1), the Genome Sequencing Program (GSP) of the National Human Genome Research Institute (https://www.genome.gov/Funded-Programs-Projects/NHGRI-Genome-Sequencing-Program), UK biobank (2) and All of Us (3). These large WGS/WES studies and biobanks have sequenced hundreds of millions of coding and non-coding genetic variants across the human genome from hundreds of thousands of individuals, evaluating their relationship to diseases and traits.

Variant functional annotation represents a powerful and effective approach to leveraging functional information from many different bioinformatics sources to elucidate the multi-faceted functions of genetic variants for a wide range of analyses of WGS/WES studies and Genome-Wide Association Studies (GWAS) (4-14). A variety of functional annotations have been developed to measure multiple aspects of biological functionality of variants, including protein function (15-17), conservation (18,19), epigenetics (20,21), spatial genomics (22,23), network biology (24), mappability (25), local nucleotide diversity (26) and integrative composite annotations (4,27-29). These annotations have successfully prioritized plausible causal variants of underlying GWAS signals according to their functional impact in experimental studies following GWAS findings (5,30), localizing causal variants in fine-mapping studies (4,8), partitioning heritability in GWAS (6), predicting genetic risk (6,7,9), and improving rare variant (RV) analysis of WGS association studies (12-14,31). For example, large-scale WGS/WES studies (1,3,32,33) assess the associations between complex diseases/traits and coding and non-coding rare variants across the genome. The recently developed STAAR method incorporates multi-faceted variant functional annotations to boost the power of rare variant association tests in WGS/WES studies (12-14).

There is a pressing need to develop a comprehensive whole genome variant functional annotation database and browser for online queries to facilitate analysis and interpretation of GWAS and WGS/WES studies, as well as software that functionally annotates any GWAS and WGS/WES study for downstream statistical genetic analysis. Although there are several existing variant functional annotation databases, such as CADD (5,34), VEP (35), Annovar (36), WGSA (37), they have several limitations. First, these existing variant annotation databases have narrow scope. None of the databases provides overall and ancestry-specific allele frequencies from gnomAD (38) and TOPMed (1), and ClinVar information (39). Second, these resources have limited online query capabilities, and do not provide a user-friendly variant function annotation browser that summarizes and visualizes multi-faceted functional annotations of a single variant and multiple variants in a gene or a region. For example, WGSA does not provide a browser for querying variant functional annotations online. VEP provides a browser but only a few annotations. CADD allows for querying a single variant or variants in a region, but displays the annotation results in a large table that is difficult to navigate. Furthermore, most of these resources do not allow for gene- and region-level variant annotations. None of these databases provides a summary or visualization of query results. Third, there is a lack of scalable and easy-to-use tools that satisfy the need of functionally annotating large-scale WGS/WES studies. Existing functional annotation databases and tools are not scalable for functionally annotating a massive number of variants in large-scale WGS/WES studies. Moreover, few of the currently available functional annotation tools can provide organized output in a format that is both storage efficient and ready to be used in downstream statistical genetic analyses, such as fine-mapping (4,11), heritability (6), rare-variants association tests (13,14). There is a pressing community need to develop a convenient and comprehensive functional annotation tool that annotates any WGS study dataset at scale and generates a functionally annotated genotype file in an organized and compressed format, that can be readily integrated into the downstream analysis.

We developed Functional Annotation of Variants Online Resources (FAVOR), a comprehensive whole genome variant annotation database and a variant browser that provides hundreds of functional annotation scores for all possible 9 billion Single Nucleotide Variants (SNVs) and observed short insertions/deletions (indels) from a variety of biological functional aspects. FAVOR provides a fast, convenient, and user-friendly web interface that features online single variant and gene-/region-level variant queries. Search results are well-organized and visualized, according to their major functional categories. FAVOR distinguishes itself from the limitations of existing tools by providing functional annotation information that can be easily viewed through multiple functional category-based blocks and tables directly on its web interface. On top of that, FAVOR automatically generates dynamic summaries of search results by identifying important functional scores of the queried variant. These FAVOR unique features grant users immediate and intuitive insight into the search results while still maintaining users’ access to the comprehensive display of multi-faceted functional scores. We have provided a comparison between FAVOR with the existing annotation databases (Supplementary Table 1).

We have also developed FAVORannotator, a tool that functionally annotates the genotype data of any WGS/WES study at scale using the FAVOR database (GRCh38 build) and stores the genotype data and their aligned functional annotation data in an annotated Genomic Data Structure (aGDS) file. The proposed aGDS data format extends the Genomics Data Structure (GDS) format (40), by storing the genotype data and the corresponding functional annotation data in a single file, making downstream integrative analysis of variants with their functional annotations more efficient and convenient. The GDS format is highly storage-efficient, with a compression rate of a thousand times compared with the VCF format. FAVORannotator is scalable and computationally efficient for functionally annotating large-scale WGS/WES studies, for example, it completes the functional annotation of 1 billion variants in 38 CPU hours and storing those data in an aGDS file of size 488 GB.

## FAVOR DATABASE

The FAVOR relational functional annotation database provides comprehensive multi-faceted variant functional annotations of all possible 9 billion SNVs in the whole genome by integrating data from multiple different sources, including CADD v1.5 (5,34), GENCODE v31(41), Annovar (36), WGSA (37), ClinVar (39), ENCODE (42), SnpEff (43), 1000 Genome (44), TOPMed Bravo Freeze 8 (1), gnomAD v3(38), and other individual studies (25,28,45-48). The preprocessing stage assigns the functional annotation values to each variant by using the variant as the primary key of the relational database. The FAVOR database is built using the PostgreSQL relational DBMS (Database Management System) for storing and retrieving variant annotation data and using a multi-table design that supports efficient integration of different types of scores. Specifically, it stores 160 functional annotation values for all possible 8,892,915,237 SNVs, and 79,997,898 observed indels from TOPMed Freeze 8 in 20 TB of space (Supplementary Table 2). These functional annotations are organized into 12 major types, including Variant Category, Allele Frequencies (AFs), ClinVar, Integrative Scores, Protein Functions, Conservation, Epigenetics, Chromatin States, Local Nucleotide Diversity, Mutation Density, Mappability and Proximity (Supplementary Table 3). The FAVOR database can be downloaded from the FAVOR website.

## FAVOR ONLINE PORTAL

The online FAVOR portal facilitates fast and convenient online functional annotation query using an R shiny app (Figure 1). It allows users to search for a single variant (either in position format or rsID), multiple variants in a gene or genomic region (either in position format or gene name), or batches of tens of thousands of variants. The variant functional annotation results are displayed in tabular overviews in a summary tab (Figure 2), a full tables tab (Figure 3), and visualized using histograms (Figure 4).

**Figure 1.**
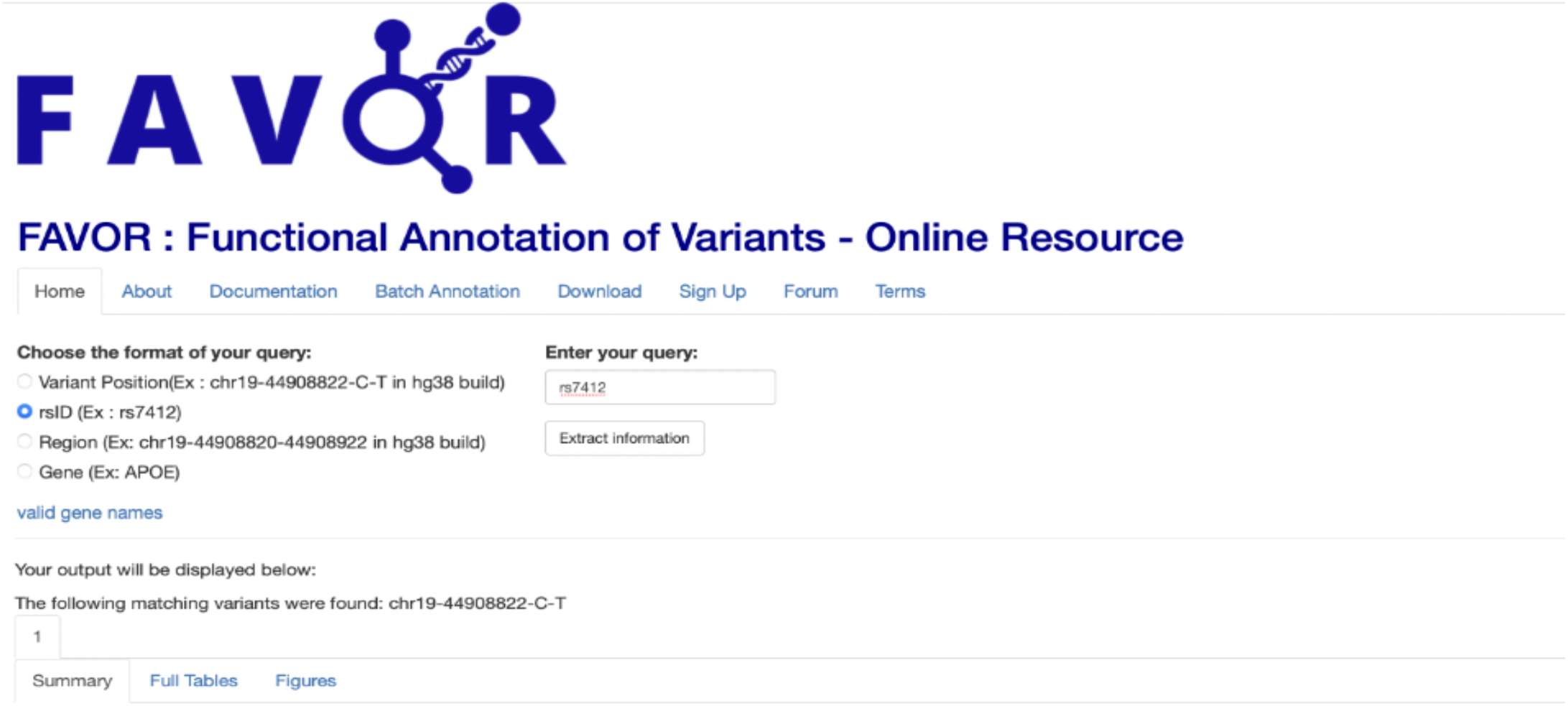
FAVOR web interface: This online portal provides a convenient web interface allowing for variant, gene, and region-level variant annotation queries. The home page displays the supported query methods, and examples of expected input.

**Figure 2.**
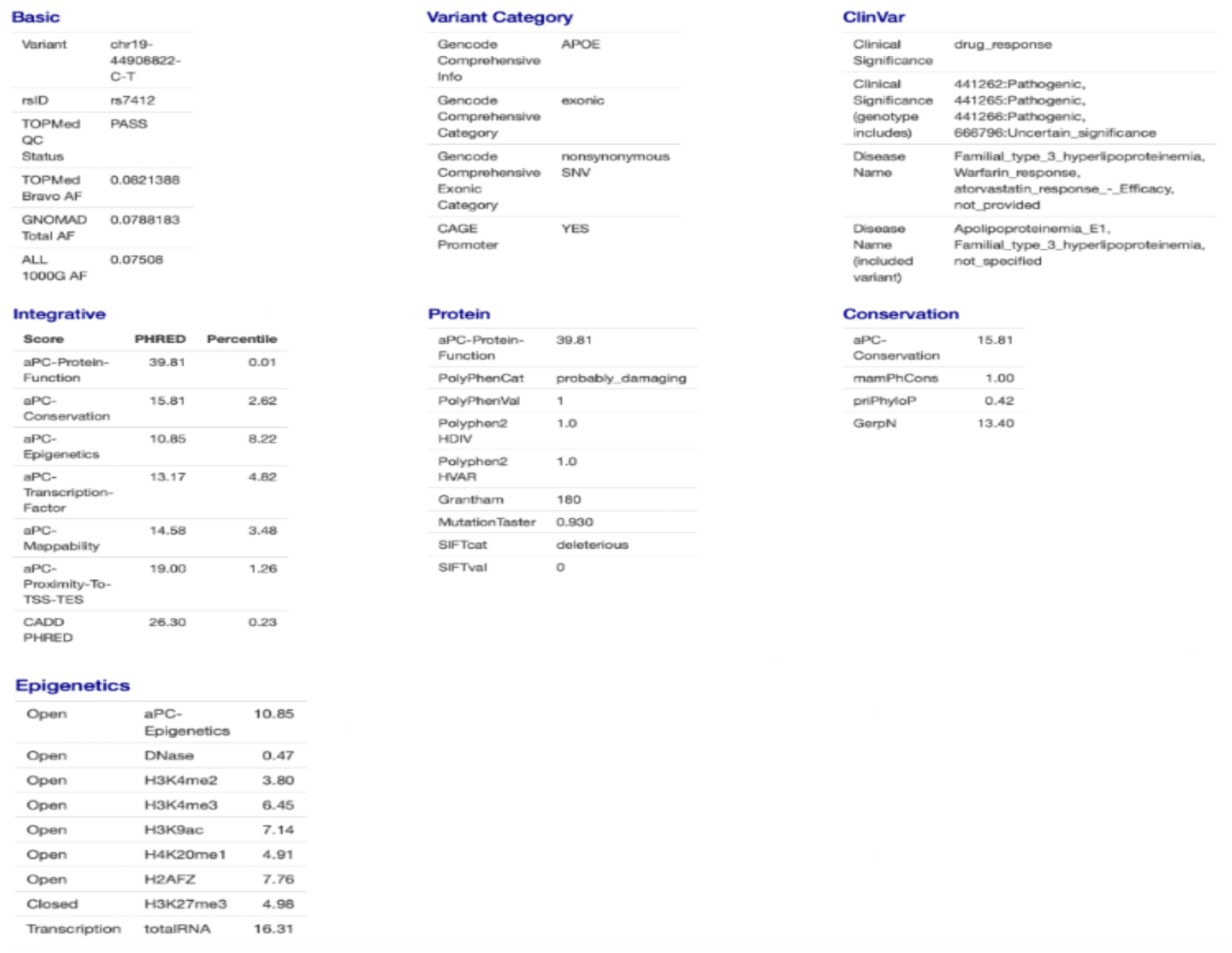
Single Variant Query Summary page: The Single Variant Query Summary page shows a dynamic overview of the filtered annotations with evidence for plausible functional consequences. For example, the annotations are displayed if Polyphen scores equal to 1 (probably_damaging), SIFT score equals to 0 (deleterious), ClinVar Significance is Pathogenic, and the integrative scores that are greater than 10 on the PHRED scale.

**Figure 3.**
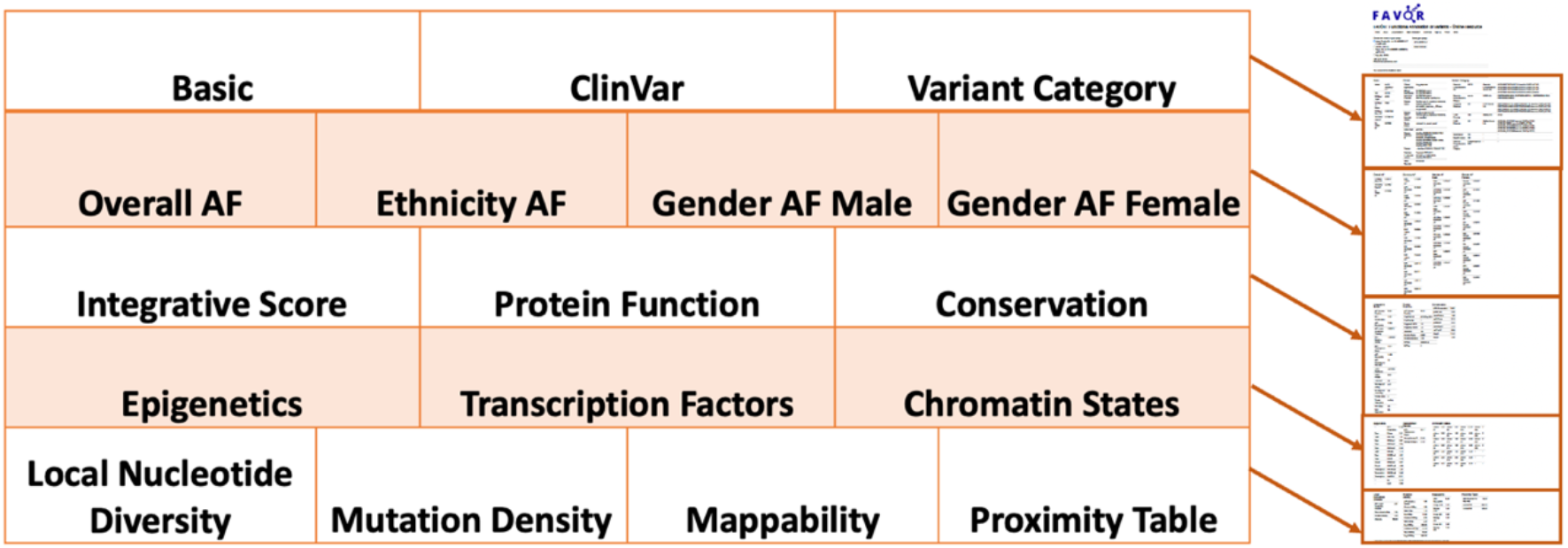
Single Variant Query Functional Annotation Tabulation: The Full Tables tab in the FAVOR Single Variant Online Query organizes functional annotation results in blocks defined by annotation types.

**Figure 4.**
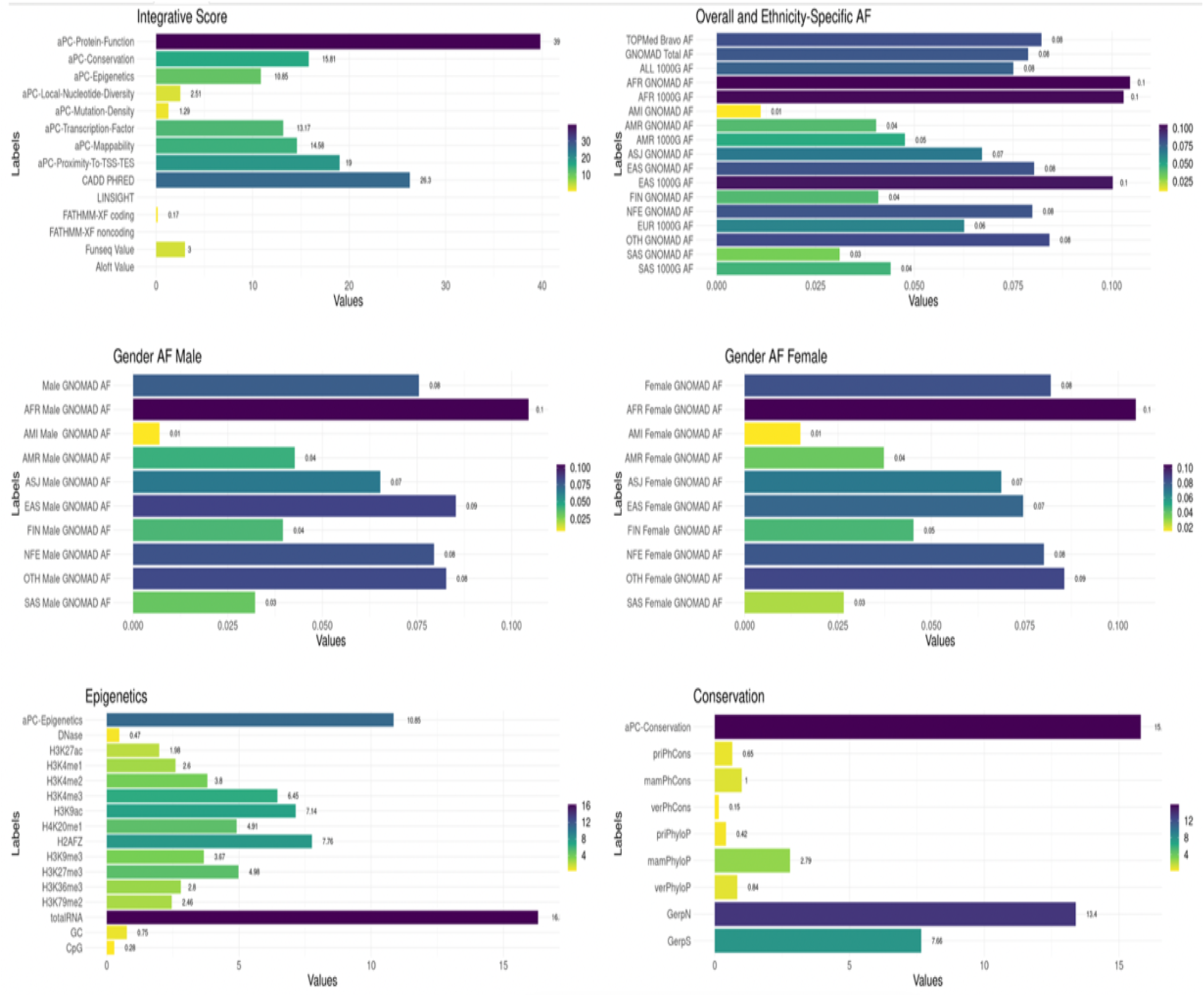
Single Variant Query Functional Annotation Visualization: The Figures tab in the FAVOR Single Variant Online Query displays a visualization of the functional annotation results of a queried variant in histogram.

The FAVOR web interface is exceptionally nimble. Single Variant Search (both variant position and rsID) renders results on the webpage immediately, while Gene-based and Region-based Variant Search takes just a few seconds to display results, and Batch Annotation directly generates the annotation results for up to 10,000 variants allowing for a range of input file formats. This fast response speed is the product of its backend database indices and table design. The indices employ a diverse set of data structures, each tailored toward specific functionalities. The table design relies upon an original primary key (a combined string that consists of variant chromosome position and reference and alternative allele, e.g., 19-44908822-C-T) that efficiently relates the tables with regard to both computation and storage. This implementation enables the fast query of 160 annotations for all 9 billion SNVs at the variant, gene and region levels.

### Single Variant Search

For Single Variant Search, users can input a variant position (in hg38 build) or an rsID. The retrieved functional annotation results are displayed in three tabs: Summary, Full Table, Figures. The Summary Tab gives an overview of the biological functionality of a variant by providing the filtered annotations that flag the variant as plausibly functional, for example, Polyphen scores equal to 1 (probably_damaging), SIFT score equals to 0 (deleterious), or ClinVar Significance is Pathogenic, and the integrative scores greater than 10 on the PHRED scale (in the top 10% of the genome). By selecting and presenting the most informative functional annotation of a queried variant in the summary tab avoids overwhelming users with a large amount of information.

The Full Tables tab displays all functional annotation scores - organized into 17 blocks of annotation groups (Figure 4). These blocks are Basic, ClinVar, Variant Category, Overall Allele Frequencies (AFs), Ethnicity-Specific AF, Gender AF, Integrative Score, Protein Function, Conservation, Epigenetics, Transcription Factors, Chromatin States, Local Nucleotide Diversity, Mutation Density, Mapability, Proximity Table.

Different groups of functional annotation depict the variants from multiple functional perspectives. For example, ClinVar reports the relationships between genetic variants and phenotypes (39). FAVOR provides critical information from ClinVar, including Clinical Significance, Disease Name, Review Status, Disease Database ID, and Gene Reported related to the variants. Variant Category annotations provide the consequences of the genetic variants in the context of gene, categorical regulatory information, and the relative location of the variant with the closest gene (Supplementary Table 3).

FAVOR integrates multiple AFs of observed variants from multiple variant databases, including the overall AF, ethnicity-specific AF and gender-specific AF from 1000 Genome (44), TOPMed Bravo Freeze 8 (1) and gnomAD v3 (38). FAVOR provides multiple integrative scores for both coding and non-coding variants, including CADD v1.5 (5,34), LINSIGHT (49), FATHMM-XF (29), FunSeq (46), Aloft (28) and annotation Principal Components (aPCs). The aPCs summarize multiple aspects of variant function by calculating the first variant-specific PC from the individual functional annotation scores in a functional category (12). For example, aPC-conservation is the first PC of the eight individual standardized conservation scores.

Furthermore, FAVOR displays category-specific individual functional annotations that represent multiple biological functionalities of each variant in a given functional category (Supplementary Table 3). For example, protein function scores describe various impact scores of the variant’s damages to protein function. Conservation scores summarize the conservation functional annotation of the variants (both within and between species). Epigenetics scores summarize the signals of the open chromatin markers, close chromatin markers, and transcription markers. FAVOR also provides individual annotation scores of local nucleotide diversity, mutation density, mappability (e.g., using the unconverted genome Umap and the bisulfite-converted genome Bismap) (Supplementary Table 3). Data visualization is available using histograms in the Figures tab for a more intuitive look into functional importance (Figure. 4).

### Region/Gene-based Search

For Region-/Gene-based search, users input either a gene name (official symbol), or region (starting and ending positions using the hg38 build). FAVOR will instantaneously output the functional annotation summary results of the variants in the gene or the region, as well as variant-specific annotations in a range of annotation categories. The fast display of the retrieved results of the Region/Gene-based Search is enabled through indexing and efficient multi-table database management.

The Region-/Gene-based Search summary tab provides the summary statistics of the variants in a region or a gene using several key summary tables and histograms, including Allele Frequency Distribution, GENCODE Category, ClinVar Clinical Significance, Functional Consequences and High Integrative Functional Scores (Figure 5). In the Region/Gene-based individual variant annotation table, 32 commonly used annotations (Supplementary Table 4) are displayed for each variant in the gene/region. The variants can be sorted by their values in any column. It also has a convenient search feature that allows users to filter the variants in the region/gene based on specified features and key words. For example, typing “pathogenic” in the search box above the displaying table provides only the pathogenic variants of the region/gene.

**Figure 5.**
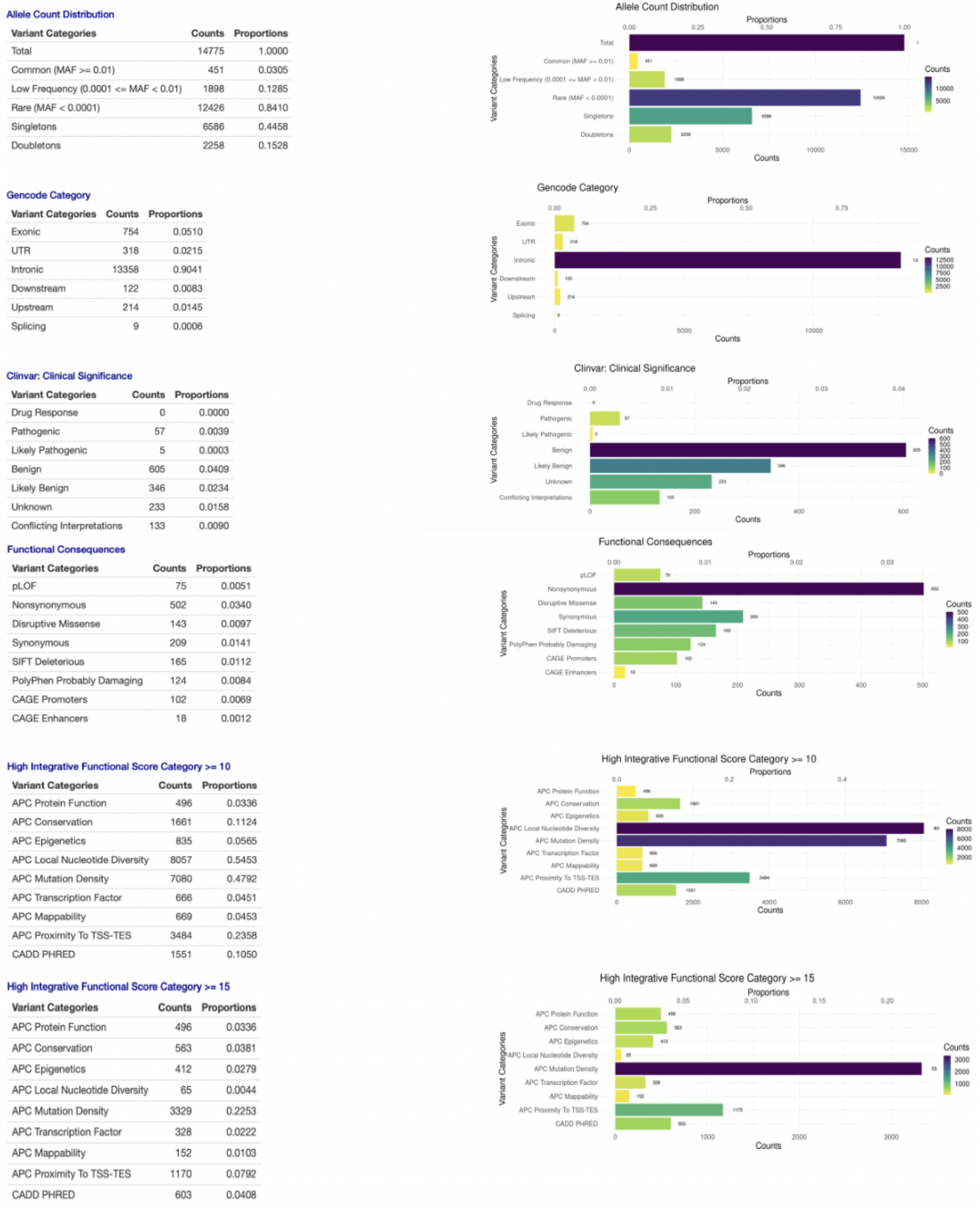
Region-/Gene-based Query Summary tab: The Summary tab of the Region-/Gene-based Query shows the multi-faceted functional annotation summary statistics of the variants in a gene or a region.

### Batch Annotation

Batch Annotation provides functional annotations of a list of variants submitted by users in a file. It supports multiple file formats as input, including CSV, TSV, VCF, XLS, and RDS. Multiple formats and IDs of variants are also supported. For example, each row of a text file can specify a variant’s chromosome, position, reference, and alternative allele value (e.g., 1-10253-CTA-C), or a variant’s chromosome and position values (e.g., 1-10253), or an rsID (rs868413313). Users can upload the variants list using the above file formats on the FAVOR batch annotation page. Batch annotation files are currently limited to 10,000 variants in the interest of online wait time. It takes less than less than 1 minute to annotate 1,000 variants. The annotation results containing 160 annotations of the variants in the submitted variant list are available for download. FAVORannotator, discussed below, can be used to handle functional annotations of a larger number of variants, e.g., hundreds of millions of variants in a WGS/WES study.

## ANNOTATED GENOMIC DATA STRUCTURE (aGDS)

Variant Call Format (VCF) (50) has been frequently used for storing variant call data of sequencing studies. However, VCF is text-based and thus inefficient with regard to storage, particularly for large-scale WGS data of hundreds of thousands to millions of subjects that have hundreds of millions to billions of variants. The recently developed Genomic Data Structure (GDS) format (40) provides a storage-efficient format to store WGS data. The GDS format has a compression rate of 1000 times compared to the VCF format. However, it does not incorporate variant functional annotations. We developed the annotated Genomic Data Structure (aGDS) format (Figure 6), that extends the GDS format by integrating both genotypes in a WGS study and variant functional annotations in a single file. There are three main advantages of the aGDS format. First, it provides fast query and simultaneous retrieval of genotype and matched functional annotation data defined by flexible filtering criteria. Second, it is convenient to integrate an aGDS file into functionally informed downstream analysis pipelines, such as STAARpipeline for rare variant association analysis (13). Third, it is also highly storage-efficient for genotype and functional annotation data. An aGDS file containing TOPMed Freeze 8 WGS data, including both genotype and functional annotations of 140,306 samples, only takes 487 GB, that is three orders of magnitude smaller compared to the VCF files (Supplementary Table 2).

**Figure 6.**
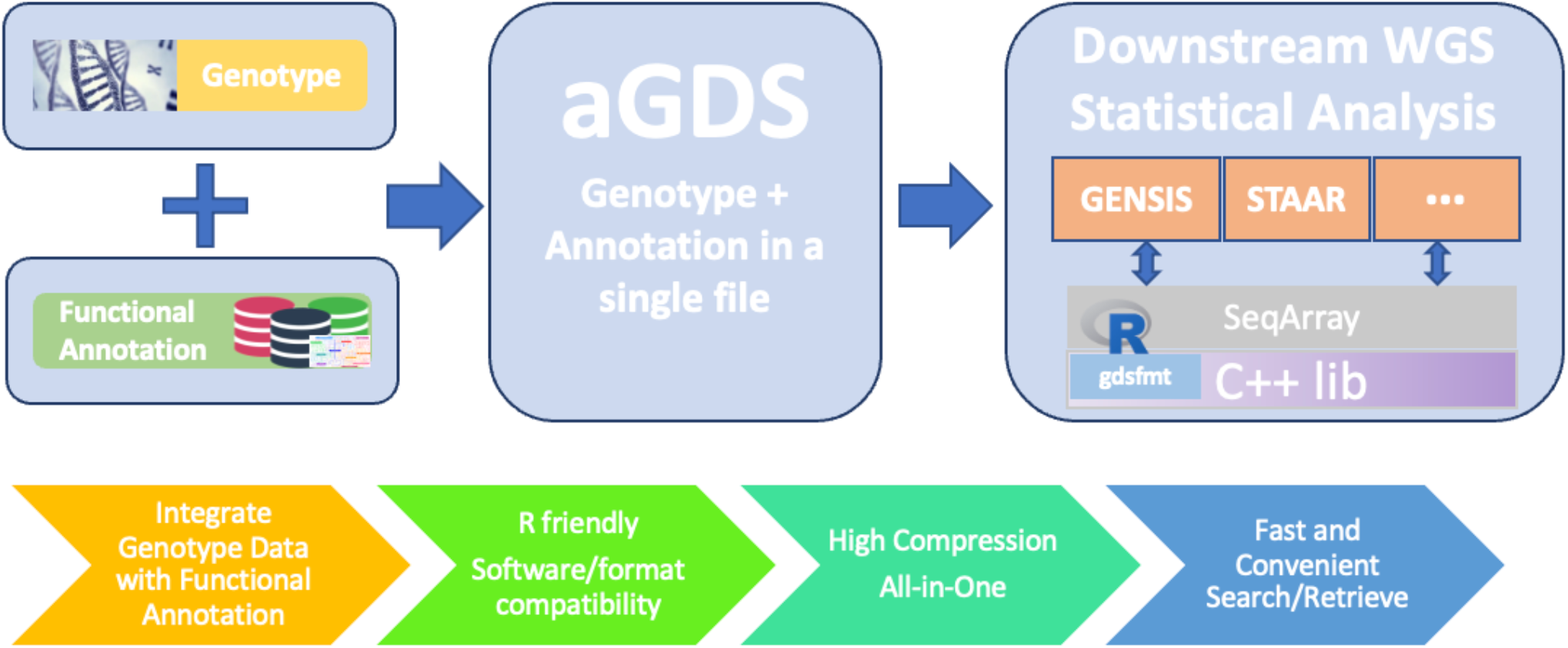
Features of the annotation Genomics Data Structure (aGDS) format: This figure shows the features of the aGDS format and the process of creating aGDS files by combining functional annotations with genotype data.

## FAVORANNOTATOR

FAVORannotator is an open-source tool that uses the FAVOR database to functionally annotate and efficiently store genotype and variant functional annotation data of a WGS/WES study in an aGDS file, and to facilitate downstream association analysis (Figure 7). FAVORannotator only requires genotype data or a variant list as input and automatically annotates the genotype data or the variant list, generating an aGDS file as an output. The former facilitates rare variant association analysis using individual level data, while the latter facilitates rare variant meta-analysis using summary statistics, by incorporating functional annotations, e.g., using STAAR (13).

**Figure 7.**
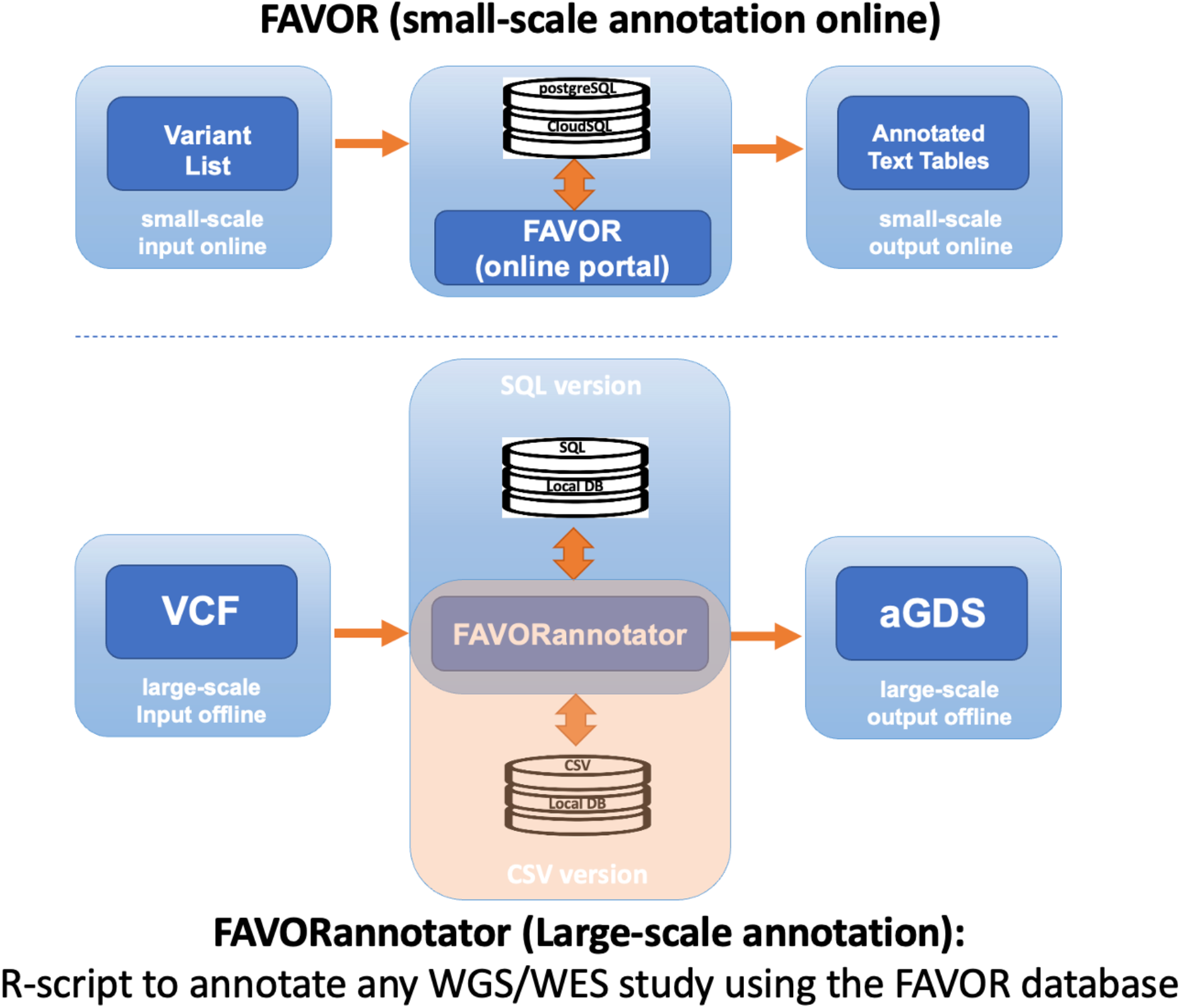
Graphical Representation of the features of FAVOR batch annotation and FAVORannotator: For small-scale annotation (up to 10,000 variants), batch annotation can be used for online annotation at the FAVOR website. For large-scale annotation, e.g., hundreds of millions of variants in a Whole Genome/Exome Sequencing (WGS/WES) study, FAVORannotator can be used for annotation in a local cluster or a cloud platform, e.g., Amazon Web Services (AWS) and Google Cloud Platform (GCP). FAVORannotator uses the backend database FAVOR, which is available in the SQL or CVS formats, and outputs an aGDS file that integrates genotype and annotation data in a single file.

Time and memory resources for annotating a large number of variants using FAVORannotator are very attractive, especially for large-scale WGS/WES datasets, such as TOPMed, GSP and UK Biobank. For example, FAVORannotator produces an annotated genotype file in the aGDS format for n=180,000 whole genome samples with 900 million variants of the TOPMed Freeze 10a WGS data in 38 hours, and for n=60545 whole genome samples of 450 million variants of GSP-CCDG Freeze 2 WGS data within 30 CPU hours. FAVORannotator has also been implemented as a workflow in the cloud-based platforms, including DNAnexus (UK Biobank), AnVIL (NHGRI) and BioData Catalyst (NHLBI) (Figure 8) (51). FAVORannotator’s efficiency keeps cloud computing costs low, for example, costing ~$25 to annotate the TOPMed Freeze 10a WGS data by chromosome in parallel, e.g., 3 CPU hours for chromosome 1.

**Figure 8.**
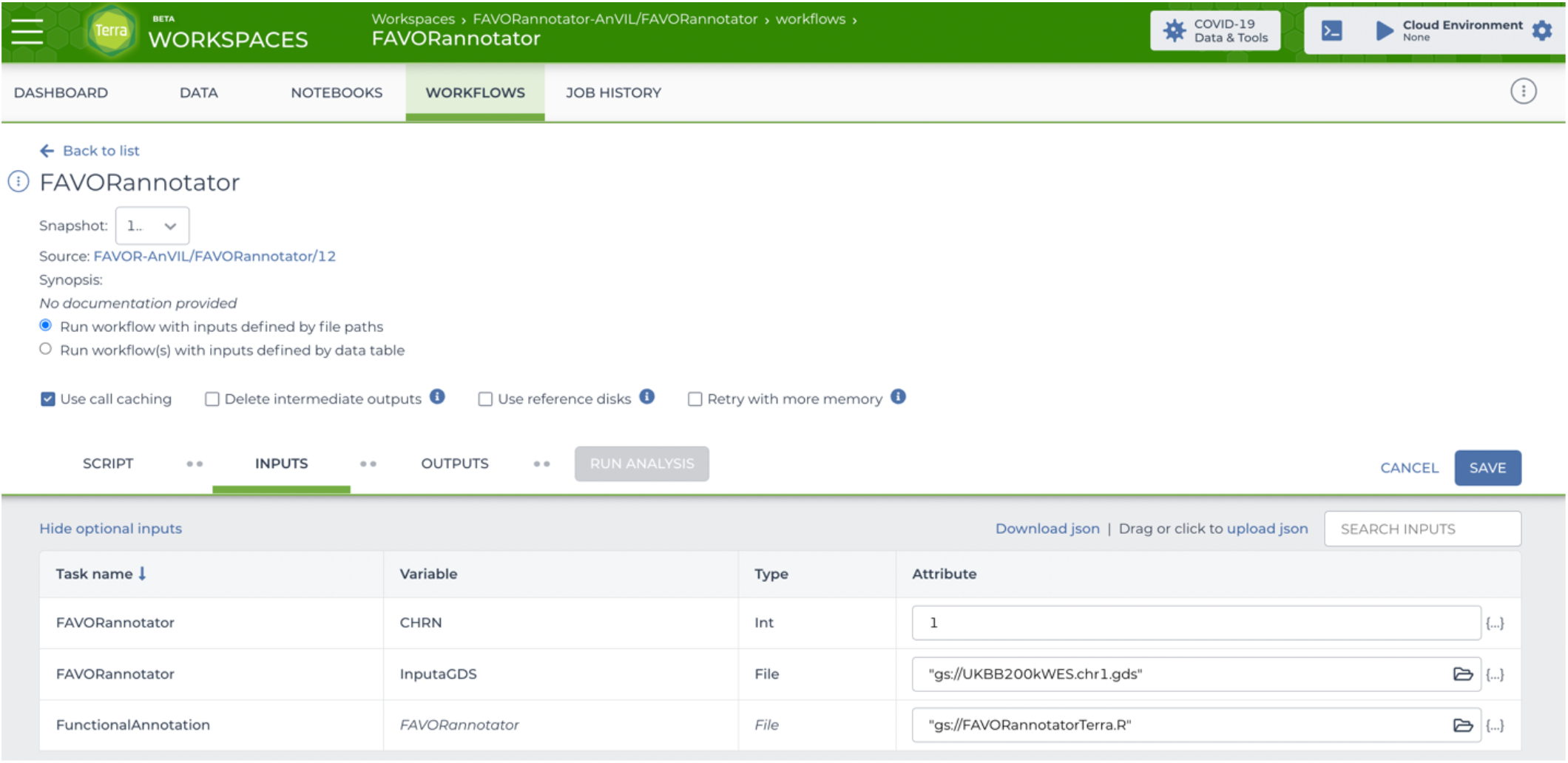
Cloud-Native FAVORannotator Workflow. The interface of the FAVORannotator Workflow on Terra.bio.

## DISCUSSION

FAVOR offers a comprehensive solution for the application of whole genome variant functional annotations, including an open access and downloadable database, a user-friendly browser, and a tool FAVORannotator, to annotate large-scale genetic data. The FAVOR database is a large relational data structure of multi-faceted functional annotations of all possible 8,812,917,339 SNVs and 79,997,898 observed indels in the human genome. It is built using a storage-efficient postgreSQL database with indexed and relational tables, that provide fast query speeds. The FAVOR web interface provides fast variant, gene, region level online multi-faceted functional annotations, as well as batch annotation. It emphasizes responsiveness while providing dynamic display and visualization features, and uses combined approaches, including visualizations, block organizations by categories, and convenient search and sorting functions, to provide a fast and convenient summary of the major functional impact of variants.

The FAVORannotator software enables researchers to use the FAVOR database to efficiently functionally annotate a genetic study at scale, such as GWAS, WGS and WES studies, and build a highly compressed and well organized aGDS file, that includes both genotype data and their annotations and can be easily integrated into downstream analysis pipelines, Together, FAVOR and FAVORannotator provide a critical infrastructure for facilitating downstream analysis and interpretation of GWAS/WGS/WES studies. Although several compression methods are available for storing WGS data, such as gzip (vcf.gz), Bgzip or BCF(52), they are subject to two major limitations. First, they are not efficient for storing large-scale WGS data. Second, they are difficult to read while compressed. For instance, although the BCF format is more storage-efficient than the VCF format, the compression rate is 100 times. In contrast, the GDS format has a compression rate of 1000 times. Furthermore, both VCF and BCF formats only store genotype data and do not store variant annotations. The aGDS format resolves both limitations successfully.

In summary, FAVOR and FAVORannotator provide an intuitive and indispensable infrastructure for facilitating downstream analysis and result interpretation of large-scale WES/WGS studies. FAVOR currently provides non-tissue specific epigenetic functional annotations for non-coding variants. It is of future interest to integrate tissue and cell-type specific epigenetic functional annotations in FAVOR. As functional annotations continue to grow in depth and breadth, we will continue to improve and expand FAVOR by integrating more and state-of-art annotations and supporting more analytical scenarios.

## AVAILABILITY

FAVORannotator is an open-source annotation tool available in the GitHub repository (https://github.com/zhouhufeng/FAVORannotator)

The FAVOR essential database (containing 20 essential functional annotation scores) for all possible SNVs (8,812,917,339) and observed Indels (79,997,898) in Build GRCh38/hg38 is hosted on Harvard Dataverse (https://doi.org/10.7910/DVN/1VGTJI).

The FAVOR full database (containing 160 essential functional annotation scores) for all possible SNVs (8,812,917,339) and observed Indels (79,997,898) in Build GRCh38/hg38 is hosted on Harvard Dataverse (https://doi.org/10.7910/DVN/KFUBKG).

