## Supplementary for "FAVOR: Functional Annotation of Variants Online Resource and Annotator for Variation across the Human Genome"

### Supplementary Material

#### Supplementary Tables

**Supplementary Table 1**. Comparison of FAVOR with other functional annotation tools. This figure compares FAVOR with existing functional annotation databases in several aspects: (1) whether a web interface is implemented; (2) whether functional annotation results are organized in functional blocks or categories; (3) whether visualization of queried results is provided on the browser; (4) whether online and offline annotation tool of a set of variants is available; (5) whether the database contains all possible 9 billion SNVs; (6) whether the annotation database supports large-scale input files with more than 500 million variants for annotation; (7) whether the functional annotation data can be easily and conveniently integrated into downstream analysis; (8) whether the functional annotation results are stored and organized alongside with their matched variant and genotype data; (9) whether cloud-native workflows are available on popular cloud platforms, e.g. Terra, DNAnexus.

| | **Web Interface** | **Organize Comprehensive Annotations in Functional Blocks for Display and Navigation** | **Results Visualization on Web Portal** | **Online and Offline Tool to annotate GWAS, WGS/WES studies** | **Annotation of all Possible 9 billion SNVs** | **Large Input Support** | **Downstream Analysis Integration** | **Annotation Output with genotype** | **Cloud-Native Workflow** |
|---|---|---|---|---|---|---|---|---|---|
| **FAVOR** | V | V | V | V | V | V | V | V | V |
| **Annovar** | V | | | | V | | | | |
| **VEP** | V | | V | V | V | | | | |
| **CADD** | V | | | V | | | | | |
| **SnpEff** | | | V | V | V | | | | |
| **WGSA** | | | | V | V | V | | | |
| **Bravo** | V | | V | | | | | | |
| **gnomAD** | V | | V | | | | | | |

**Supplementary Table 2**. File size comparison of VCF, GDS and aGDS formats: A comparison is provided for the file sizes of the aGDS format including sample-level genotypes with the VCF and GDS formats using the Freeze 2 Whole Genome Sequencing (WGS) data of the Centers for Common Disease Genomics (CCDGs) of the NHGRI Genome Sequencing Program (n=60,545) and the Freeze 8 data of the NHLBI Trans-Omics for Precision Medicine Program (TOPMed) (n=140,306). The results show that the aGDS file format has a much higher compression rate and a much smaller file size compared to the VCF file format.

| **Dataset** | **Sample Size** | **VCF Size** | **GDS Size** | **Annotation in Text Table** | **Annotation only aGDS size** | **Total (genotype + annotation) aGDS size** |
|---|---|---|---|---|---|---|
| CCDG Freeze 2 | 60,545 | 97 TB (100%) | 355 GB (0.366%) | 4 TB (100%) | 31 GB (0.775%) | 386 GB (0.382%) |
| TOPMed Freeze 8 | 140,306 | 456.7 TB (100%) | 378 GB (0.083%) | 20 TB (100%) | 100 GB (0.50%) | 478 GB (0.10%) |

**Supplementary Table 3**. Detailed functional annotations of each block in FAVOR: This table provides a list of detailed functional annotation channels of each functional block in FAVOR. Abbreviations: AF = alternate allele frequency, UCSC = University of California Santa Cruz, AFR = African/African American, AMR = Native American, EAS = East Asian, EUR = European, NFE = non-Finnish European, FIN = Finnish, SAS = South Asian, AMI = Latino/Admixed American, ASJ = Ashkenazi Jewish, OTH = other, aPC = annotation principal component, Cat=category, Val=value, pri=prime, mam=mammal, ver=vertebrate,

| Block | Detail Functional Annotation |
|---|---|
| Basic | Variant, rsID, TOPMed QC Status, TOPMed Bravo AF, gnomAD Total AF, ALL 1000G AF |
| ClinVar | Clinical Significance, Clinical Significance (genotype includes), Disease |

| | |
|---|---|
| | Name, Disease Name (included variant), Review Status, Allele Origin, Disease Database ID, Disease Database ID (included variant), Gene Reported |
| Variant Category | Gencode Comprehensive Info, Gencode Comprehensive Category, Gencode Comprehensive Exonic Category, Gencode Comprehensive Exonic Info, UCSC Info, UCSC Exonic Info, RefSeq Info, RefSeq Exonic Info, Disruptive Missense, CAGE Promoter, CAGE Enhancer, GeneHancer, SuperEnhancer |
| Overall AF | TOPMed Bravo AF, GNOMAD Total AF, ALL 1000G AF |
| Ethnicity AF | AFR 1000G AF, AFR GNOMAD AF, AMR 1000G AF, AMR GNOMAD AF, EAS 1000G AF, EAS GNOMAD AF, EUR 1000G AF, NFE GNOMAD AF, FIN GNOMAD AF, SAS 1000G AF, SAS GNOMAD AF, AMI GNOMAD AF, ASJ GNOMAD AF, OTH GNOMAD AF |
| Gender AF Male | Male GNOMAD AF, AFR Male GNOMAD AF, AMI Male GNOMAD AF, AMR Male GNOMAD AF, ASJ Male GNOMAD AF, EAS Male GNOMAD AF, FIN Male GNOMAD AF, NFE Male GNOMAD AF, OTH Male GNOMAD AF, SAS Male GNOMAD AF |
| Gender AF Female | Female GNOMAD AF, AFR Female GNOMAD AF, AMI Female GNOMAD AF, AMR Female GNOMAD AF, ASJ Female GNOMAD AF, EAS Female GNOMAD AF, FIN Female GNOMAD AF, NFE Female GNOMAD AF, OTH Female GNOMAD AF, SAS Female GNOMAD AF |
| Integrative Score | aPC-Protein-Function, aPC-Conservation, aPC-Epigenetics-Active, aPC-Epigenetics-Repressed, aPC-Epigenetics-Transcription, aPC-Local-Nucleotide-Diversity, aPC-Mutation-Density, aPC-Transcription-Factor , aPC-Mappability, aPC-Proximity-To-TSS-TES, CADD RawScore, CADD PHRED, LINSIGHT, FATHMM-XF, Funseq Value, Funseq Description, Aloft Value, Aloft Description |
| Protein Function | aPC-Protein-Function, PolyPhenCat, PolyPhenVal, Polyphen2 HDIV, Polyphen2 HVAR, Grantham, MutationTaster, MutationAssessor, SIFTcat, SIFTval |
| Conservation | aPC-Conservation, priPhCons, mamPhCons, verPhCons, priPhyloP, mamPhyloP, verPhyloP, GerpN, GerpS |
| Epigenetics | aPC-Epigenetics-Active, aPC-Epigenetics-Repressed, aPC-Epigenetics- |

| | |
|---|---|
| | Transcription, DNase, H3K27ac, H3K4me1, H3K4me2, H3K4me3, H3K9ac, H4K20me1, H2AFZ, H3K9me3, H3K27me3, H3K36me3, H3K79me2, totalRNA, GC, CpG |
| Transcription Factors | aPC-Transcription-Factor, RemapOverlapTF, RemapOverlapCL |
| Chromatin States | cHmm E1, cHmm E2, cHmm E3, cHmm E4, cHmm E5, cHmm E6, cHmm E7, cHmm E8, cHmm E9, cHmm E10, cHmm E11, cHmm E12, cHmm E13, cHmm E14, cHmm E15, cHmm E16, cHmm E17, cHmm E18, cHmm E19, cHmm E20, cHmm E21, cHmm E22, cHmm E23, cHmm E24, cHmm E25 |
| Local Nucleotide Diversity | aPC-Local-Nucleotide-Diversity, RecombinationRate, NuclearDiversity, bStatistic |
| Mutation Density | aPC-Mutation-Density, Common100bp, Rare100bp, Sngl100bp, Common1000bp, Rare1000bp, Sngl1000bp, Common10000bp, Rare10000bp, Sngl10000bp |
| Mappability | aPC-Mappability, Umap k100, Bismap k100, Umap k50, Bismap k50, Umap k24, Bismap k24 |
| Proximity Table | aPC-Proximity-To-TSS-TES, minDistTSS, minDistTSE |

**Supplementary Table 4.** List of functional annotations in the Full Table tab of FAVOR gene-/region-based query. Abbreviations: AF = alternate allele frequency, UCSC = University of California Santa Cruz, AFR = African/African American, AMR = Native American, EAS = East Asian, EUR = European, NFE = non-Finnish European, FIN = Finnish, SAS = South Asian, AMI = Latino/Admixed American, ASJ = Ashkenazi Jewish, OTH = other, aPC = annotation principal component, Cat=category, Val=value.

| **Category** | **Detail Functional Annotation** |
|---|---|
| Variant Details | rsID, TOPMed QC Status |

| | |
|---|---|
| Allele Frequency | TOPMed Bravo AF, Total GNOMAD AF, ALL 1000G AF, EUR 1000G AF, AFR 1000G AF, AMR 1000G AF, SAS 1000G AF |
| | Genecode Comprehensive Info, Gene Reported, Genecode Comprehensive Category, Genecode Comprehensive Exonic Category |
| Clinical Annotation | Clinical Significance, Clinical Significance (genotype includes), Disease Name, Disease Name (included variant), Review Status, Allele Origin, Disease Database ID, Disease Database ID (included variant) |
| Protein Function | SIFTcat |
| Epigenetics | CAGE Promoter, CAGE Enhancer, GeneHancer |
| Integrative Scores | Fathmm XF, aPC-Epigenetics-Active, aPC-Epigenetics-Repressed,aPC-Conservation, aPC-Protein-Function, aPC-Local-Nucleotide-Diversity, aPC-Mutation-Density |
